# Generation of virtual populations for quantitative systems pharmacology through advanced sampling methods

**DOI:** 10.1101/2025.06.24.661262

**Authors:** Miriam Schirru, Tristan Brier, Maxime Petit, Didier Zugaj, Pierre-Olivier Tremblay, Fahima Nekka

**Affiliations:** Laboratoire de recherche en pharmacométrie, Faculté de pharmacie, Université de Montréal, Montréal, Canada; Syneos Health, Clinical Pharmacology, Québec, Canada; Centre de recherches mathématiques, Université de Montréal, Montréal, Canada

**Keywords:** Quantitative systems pharmacology, virtual patient, DREAM toolbox, mathematical modeling, differential evolution Markov chain Monte Carlo

## Abstract

Virtual population (VPop) generation is a central component of quantitative systems pharmacology (QSP), involving the sampling of parameter sets that represent physiologically plausible patients and capture observed inter-individual variability in clinical outcomes. This approach poses challenges due to the high dimensionality and often non-identifiability nature of many QSP models. In this study, we evaluate the performance of the DREAM(ZS) algorithm, a multi-chain adaptive Markov chain Monte Carlo (MCMC) method for generating VPop. Using the Van De Pas model of cholesterol metabolism as a case study, we compare DREAM(ZS) to the single-chain Metropolis-Hastings (MH) algorithm introduced by Rieger et al. Our comparison focuses on convergence behavior, parametric diversity, and posterior coverage, in relation to the ability of each method to explore complex parameter distributions and maintain correlations. DREAM(ZS) demonstrates superior exploration of the parameter space, reducing boundary accumulation effects common in traditional MH sampling, and restoring parameter correlation structures. These advantages are attributed in part to its adaptive proposal mechanism and the use of a bias-corrected likelihood formulation, which together contribute to a better parameters space sampling without compromising model fit. Our findings contribute to the ongoing development of efficient sampling methodologies for high-dimensional biological models, introducing a promising and easy to use alternative for VPop generation in QSP, expanding the methodological approaches for in silico trial simulation.

## INTRODUCTION

In quantitative systems pharmacology (QSP), virtual populations (VPop) are essential for understanding variability in patient responses to drug treatments (Rieger et al. 2018; Cheng et al. 2022; Coto-Segura et al. 2023). Simulating patient cohorts provides insights into drug efficacy and safety across a range of clinical scenarios, supporting more informed decision-making in drug development. This ability to predict variability reduces risks, optimizes resources, and minimizes late-stage failures. VPop generation methods have been applied to diverse fields, including metabolic diseases, oncology, and immunology, showing their versatility and impact (Rieger et al. 2022; Wang et al. 2023; Braniff et al. 2024). In the context of large QSP models, VPop offer several advantages. By sampling a wide range of parameter sets that reflect biological variability, VPop allow for handling the complexity of QSP models while providing insights into diverse patient outcomes. This is valuable when individual model parameters are difficult to identify, as the virtual population calibration process constrains the parameter space to produce physiologically plausible predictions. VPop also integrate diverse data sources, including preclinical studies, clinical trials, and real-world evidence, ensuring that their predictions remain closely aligned with observed variability in clinical data (Cheng et al. 2022; Braniff et al. 2024; Chougule et al. 2024). Beyond quantitative accuracy, VP analysis enhances qualitative insights by identifying dynamical patterns across a range of plausible parameter sets, including clinical threshold effects, where small changes in key parameters (such as immune activation or drug concentration) can lead to dramatic shifts in outcomes, like disease remission or progression (Zugaj et al. 2024; Chae 2020; Zhang and Tyson 2022). These benefits make VPop an essential tool for validating and simplifying complex models without compromising their predictive performance or biological relevance. Key challenges include capturing the diversity of parameter combinations that reflect biological variability, guaranteeing alignment with clinical data, and managing proper sampling from complex, high-dimensional parameter spaces (Rieger et al. 2018; Myers et al. 2023). Addressing these challenges is critical to advancing the use of VPop.

In the past years, several methods have been developed to generate VPop. Allen et al. uses a cholesterol metabolism model as a case study and clinical data to create parameter sets within physiologically realistic ranges (Van De Pas et al. 2012; Allen et al. 2016). Their method uses simulated annealing (SA) optimization to adjust parameters and a Bayesian acceptance-rejection sampling step to match clinical data distributions. However, this approach requires high computational effort and suffers slow convergence to the target distribution. Building on this, Rieger et al. introduced the traditional Metropolis-Hastings (MH) algorithm to sample parameters space (Rieger et al. 2018). The strength of this approach lies in its use of Bayesian inference to constrain the simulated data to match the target distribution, rather than only relying on parameter ranges. By using the likelihood formalism, MH ensures that parameter sets are evaluated against clinical data. Moreover, it was demonstrated that MH performs better compared to other methods like SA, making it a practical choice for large-scale applications (Rieger et al. 2018). However, classic MH algorithms can be limited by their slow convergence (Vrugt et al. 2009) and the difficulty in handling multimodal distributions that can be encountered in biological systems. Real-world clinical data often display complex characteristics that extend beyond the assumptions of simple likelihood functions, such as the multi-dimensional Gaussian distribution, and beyond the capabilities of single-chain MH sampling.

Other strategies, such as those proposed by Sinisi et al., address the challenge of generating VPop from non-identifiable ODE models, where multiple parameter sets yield similar observable behavior (Sinisi et al. 2020). By introducing a behavioral framework based on pairwise distinguishability and completeness, their approach generates Vpop that represent diverse patient phenotypes while avoiding redundancy. This approach improves biological fidelity and robustness in in silico clinical trials. However, its computational demands limit scalability for large parameter spaces, particularly in high-dimensional models. To better exploit nonidentifiable QSP models, VPop construction can also benefit from considering how the system dynamically responds to variations in parameters, initial conditions, and both endogenous and exogenous inputs, as we recently explored (Zugaj and Nekka 2025),(submitted). Also, Gevertz et al. (2024) demonstrated that prior distribution choices can impact the heterogeneity of the population, sometimes more than the selection criteria themselves (Gevertz and Wares 2024). Despite these refinements, existing approaches still face a fundamental challenge: ensuring a high fit to the empirical data, while maximizing parameters variability.

To address these challenges, we propose the use of DREAM (Differential evolution adaptive metropolis), an advanced version of the MH algorithm originally developed for hydrology and environmental science (Vrugt 2016). DREAM introduces multi-chain sampling, adaptive convergence mechanisms, random subdimension sampling, and outlier detection to overcome the limitations of traditional MH methods. By running multiple Markov chains in parallel, DREAM efficiently explores high-dimensional and multimodal parameter spaces while maintaining diversity across chains. The chains exchange information dynamically, helping the algorithm to avoid local minima and promote broader exploration. An additional feature of DREAM is its ability to restore correlations in the target distribution by adapting its sampling process to the evolving shape of the posterior. This ensures that interdependencies between parameters are accurately captured during sampling. At the same time, exploration mechanisms introduce randomness into the search process, enabling probing less probable regions in the parameter space and preventing premature convergence.

Parameters redundancy can result in a Vpop with many individuals with same model outputs. This reflects the heterogeneity of the targeted distribution and highlights the importance of ensuring diversity to represent the full spectrum of clinically relevant behaviors. For example, variations in cholesterol production and clearance rates can lead to identical cholesterol levels. DREAM has the ability to capture such variability, generating biologically diverse and clinically relevant VPs that enable deeper insights into patient subgroups and differential drug responses. DREAM is implemented via a MATLAB toolbox that simplifies its use (Vrugt 2016). Researchers can input parameter ranges, prior distributions, and likelihood functions, and the algorithm outputs VPs. This accessibility makes DREAM a practical solution for a wide range of QSP modeling applications, such as those considered in Sové et al. 2020; Wang et al. 2021; Schirru et al. 2024.

To our knowledge, the current study represents the first application of DREAM in QSP modeling for the generation of VPop. While the case study considered here illustrates a steady-state modeling approach, we emphasize the suitability of DREAM to handle more complex cases, including time-dependent scenarios, such as tumor growth profiles or time-dependent PK/PD studies. This adaptability makes DREAM a promising tool for a broad range of pharmacological modeling problematics.

In the current study, we evaluate DREAM in the context of a cholesterol metabolism model (Van De Pas et al. 2012), comparing its performance to the implementation of the MH algorithm proposed by Rieger et al. (Rieger et al. 2018), referred to as the reference method throughout the manuscript. Extending their metrics, we assess the ability of each method to generate Ppop with suitable properties. We will detail the methodology and present the algorithm’s key features and modifications. Our results suggest that DREAM can generate plausible, biologically diverse VPs, with implications for parameter diversity. Finally, we discuss the advantages of using this approach, emphasizing its ability to preserve key properties of the parameter space.

## METHODOLOGY

### WORKFLOW

The workflow adopted in this study to construct VPop using the DREAM(ZS) algorithm is depicted in **Fig.1a**. The process is designed to efficiently explore the parameter space, evaluate plausible parameter sets, and construct VPop optimized to fit the clinical data. This workflow consists of four stages: i) the process begins by defining the model, observables, parameter ranges and distributions, and the external data used as a reference distribution of observables; ii) configuration of the DREAM(ZS) algorithm, such as the number of chains, iterations and likelihood function, are defined; iii) the sampling process is performed iteratively across multiple Markov chains, refining parameters values based on the posterior distribution (explained below); iv) sampled parameter undergo internal processing steps to extract a Ppop. At this stage, the algorithm’s performance is assessed; v) the impact of including a reselection method to refine the transition from plausible to virtual populations is explored. Detailed explanations of each stage are provided in the following sections.

**Fig. 1.**
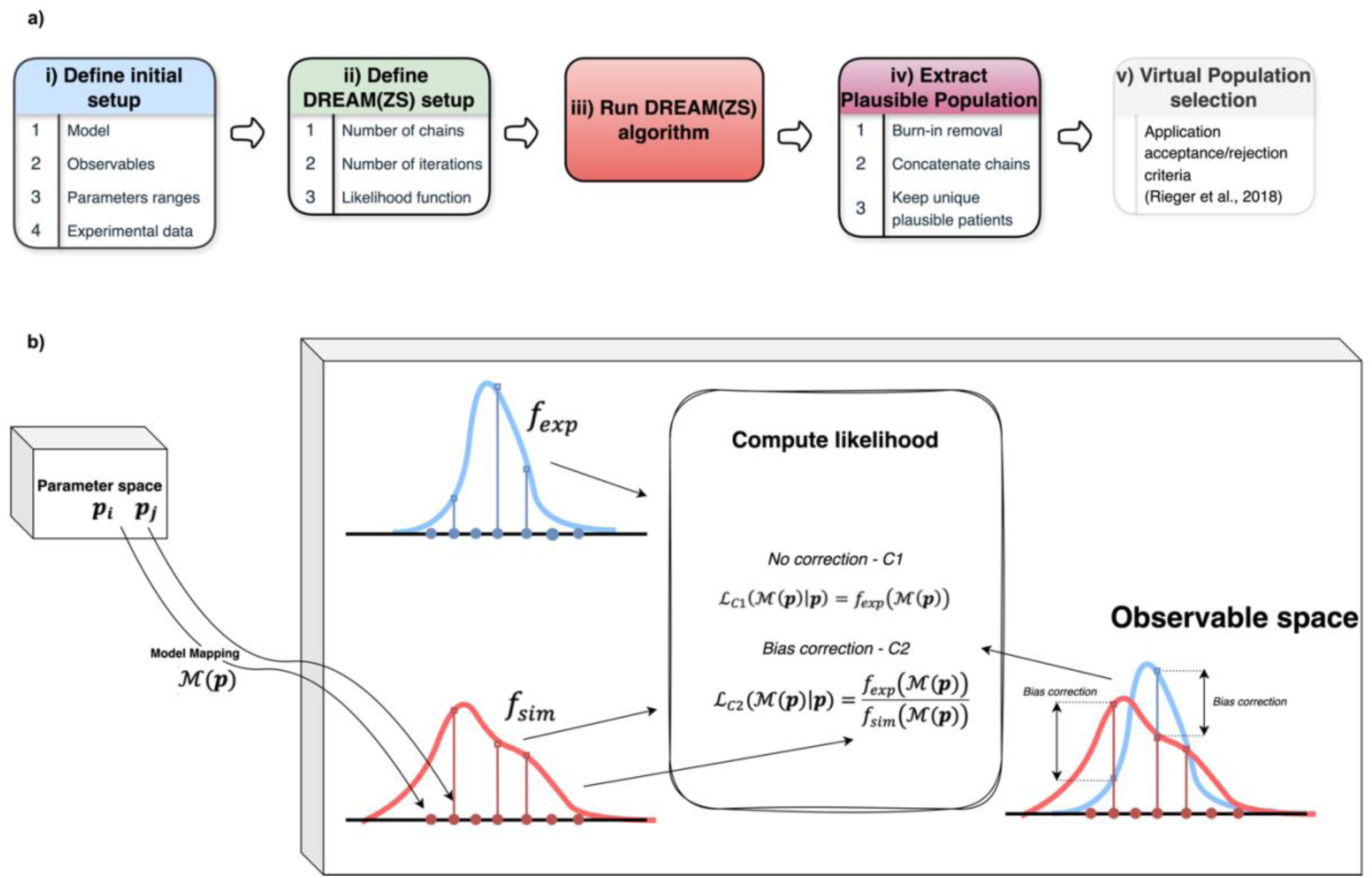
Workflow for plausible population generation and likelihood computation. a) Overview of the plausible population generation procedure. b) The parameter vector p, sampled from the parameter space (here 22 dimensional), is mapped through the model ℳ(𝒑) to generate an output in the observable space (here 3 dimensional). In the observable space, the model output is compared to the empirical probability density function 𝑓_𝑒𝑥𝑝_, which represents the distribution of experimental data. For comparison with the reference method, we use a multidimensional Gaussian PDF estimate as 𝑓_𝑒𝑥𝑝_ . Further explanations are given below. The likelihood function can be computed in two ways. 1) No correction C1, where the likelihood ℒ_𝐶1_(ℳ(𝒑)|𝒑) = 𝑓_𝑒𝑥𝑝_ (ℳ(𝒑)) is evaluated solely by the empirical density at the model output: 2) Bias correction C2, to prevent over-representation of certain regions, by the model mapping, in the observable space. In this case, a bias-corrected likelihood is used where the likelihood 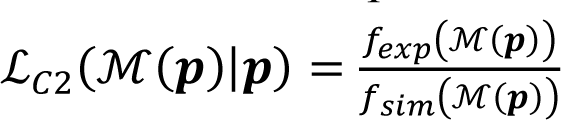 is adjusted by normalizing with the model’s own probability density estimate 𝑓_𝑠𝑖𝑚_. The right side of the figure 1b illustrates how this correction adjusts the likelihood computation by accounting for regions that are oversampled (bias correction left) or under sampled (bias correction right) by the model. The DREAM(ZS) algorithm then uses the computed likelihood to refine parameter selection.

### MODEL MAPPING INTEGRATION

The dynamical system representing the model is described using a model mapping function ℳ(𝒑, 𝒙_𝟎_, 𝒖(𝑡)) = 𝒐(𝑡) which defines how parameters 𝒑, initial conditions 𝒙_𝟎_ and inputs 𝒖(𝑡) impact the outputs 𝒐(𝑡). The outputs of this mapping can vary (steady-states values, trajectories over time, clinically relevant outcomes, AUCs, RECIST etc.).

In the current study, the model is defined as a mapping ℳ: 𝒫 → 𝒪, where 𝒫 ⊂ ℝ^𝑛^ is the parameter space (𝑛 = 22 here) , and 𝒑 ∈ 𝒫 represents a parameter vector; 𝒪 ⊂ ℝ^𝑚^ is the observable space (𝑚 = 3 here), and 𝒐 = ℳ(𝒑) ∈ 𝒪 represents the corresponding observable vector evaluated at steady states.

The probability density function of simulated observables, 𝑓_𝑠𝑖𝑚_, represents how frequently specific observable values appear under the mapping ℳ according to the prior distribution (see section below).

In the context of DREAM algorithm, model mapping is critical because it allows the evaluation of how well different parameter sets generate biologically plausible outcomes that align with empirical data. Incorporating a model mapping into DREAM introduces additional complexity, as the algorithm no longer operates solely in the parameter space but evaluates performance in the observable space. This step is necessary because we are evaluating sampled parameters based on model outputs, ensuring that only parameter sets (with real-world biological variability) are retained.

Formally, given:

- A parameter vector 𝒑 ∈ 𝒫, where 𝒫 is the biologically bounded subspace of the parameter space;

- A model mapping ℳ(𝒑) which transforms the parameter vector 𝒑 into the observable space 𝒪;

- An empirical probability density function (pdf) 𝑓_𝑒𝑥𝑝_, describing the distribution of the experimental dataset 𝐷_𝑒𝑥𝑝_ ⊂ 𝒪.

As a first case (C1), the likelihood is formulated as:

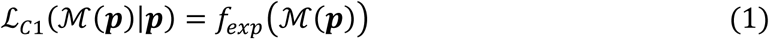

By “encoding” observation matching, this likelihood function guides the sampling of parameter vectors 𝒑 such that their corresponding outputs ℳ(𝒑) follow the distribution of 𝐷_𝑒𝑥𝑝_ . In the Bayesian framework, this alignment is formalized as the posterior distribution over the parameters 𝒑, which combines the likelihood ℒ_𝐶1_ with a prior distribution 𝑓_𝑝𝑟𝑖𝑜𝑟_(𝒑):

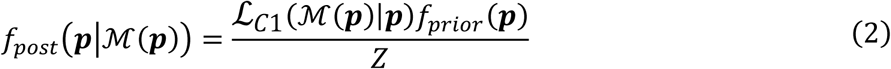

where:

- 𝑓_𝑝𝑟𝑖𝑜𝑟_(𝒑) is the prior distribution (uniform) over the parameters, including prior knowledge or constraints about plausible parameter values.

- 𝑍 is the marginal likelihood (normalization constant), defined as:

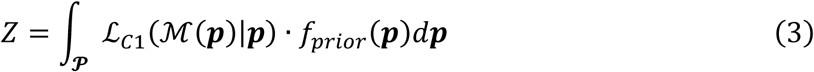

- The posterior distribution 𝑓_𝑝𝑜𝑠𝑡_(𝒑|ℳ(𝒑)) represents the updated belief about the parameters 𝒑 after incorporating 𝐷_𝑒𝑥𝑝_ .

To prevent over-representation of certain regions in the observable space 𝒪, we introduce a second case for the application of DREAM (called C2 in the following) in which the likelihood function ℒ_𝐶1_(ℳ(𝒑)|𝒑) = 𝑓_𝑒𝑥𝑝_(ℳ(𝒑)) is normalized by 𝑓_𝑠𝑖𝑚_(ℳ(𝒑)), the prior 𝑓_𝑝𝑟𝑖𝑜𝑟_(𝒑) pushed forward into the observable space by the model as follows:

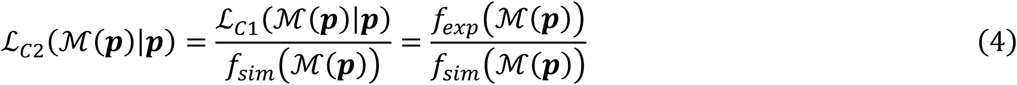

This correction ensures that the empirical density 𝑓_𝑒𝑥𝑝_ drives the posterior distribution without over-sampling certain parameter regions, as shown in **Fig.1b**. By considering the hypothesis that the data is independent from the model, a formal proof to justify this likelihood is derived in the Supplementary Material 1.1 (SM1.1.1). The construction of 𝑓_𝑠𝑖𝑚_ is detailed in the SM1.1.2.

### METROPOLIS-HASTINGS ALGORITHM IN DREAM

The MH algorithm is used to sample the parameters from the posterior distribution 𝑓_𝑝𝑜𝑠𝑡_ (𝒑|ℳ(𝒑)) and works as follows in DREAM:

1. Initialization: for each chain, start with an initial parameter vector 𝒑_𝟎_;
2. Proposal: for each chain, generate a new candidate 𝒑^∗^ from a proposal distribution 𝑞(𝒑^∗^|𝒑);
3. Acceptance ratio: compute the acceptance ratio for each proposed 𝒑^∗^

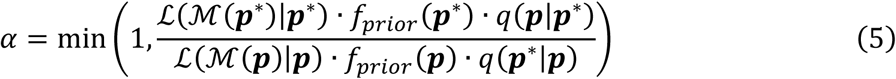

1. Acceptance step: accept 𝒑^∗^ with probability 𝛼; otherwise, keep 𝒑;
2. Iteration until convergence: repeat steps 2-4 until a predefined number of iterations is achieved (as described below).

It is worth noting that DREAM does not explicitly rely on a predefined proposal distribution 𝑞(𝒑^∗^|𝒑), as is common in standard MH or similar MCMC algorithms. Instead, the differential evolution strategy to generate proposals is used (Vrugt 2016). The likelihood computation pseudocode is detailed in SM1.2.

### THE WAY DREAM IS USED

#### Configuration

The DREAM algorithm offers different extensions to sample from the posterior distribution. The choice of the specific algorithm, DREAM(ZS), depends on computational aspects and adaptability to complex parameter spaces. DREAM(ZS) runs a minimum of three chains, each independently exploring the parameter space while interacting with others through differential evolution updates (Vrugt 2016).

In this study, the number of iterations is chosen based on a two-step process. A convergence phase (burn-in) of 5000 iterations per chain was set to allow the Markov chains to stabilize. To achieve the desired effective Ppop sample size, we set the total number of iterations to 25000 per chain, informed by the algorithm’s acceptance rate and a predefined population size. With three parallel chains, this resulted in 75000 total iterations, of which 15000 were discarded as burn-in. Convergence and acceptance rate diagnostics supporting these choices are detailed in SM1.3.1. Given a computed acceptance rate of 45%, the final Ppop size is approximately 27000 unique samples, as described in SM1.3.2. The decision to retain unique samples or allow some redundancy depends on the desired usage of the population. While some redundancy is inherent to the DREAM(ZS) algorithm and can indicate convergence toward high probability regions in the target distribution, in this study, we focus on retaining only unique parameter sets, meaning that duplicate samples are removed.

### DEFINITION OF PLAUSIBLE AND VIRTUAL POPULATION

To understand how sampled parameter sets transition from an initial exploration phase to a final selection, it is important to formally introduce the distinction between Ppop and Vpop. We first construct a Ppop, generated by sampling from a wide range of possible parameters sets, reflecting the natural variability among patients or biological systems. These parameter sets are chosen to match known biological and clinical data. While they define a Ppop, an additional refinement step is required to construct a Vpop. This step either ensures appropriate coverage of the observable space by accounting for specific constraints (Allen et al. 2016), or corrects for sampling approximations (such as assumptions about symmetry in the proposal distribution) (Rieger et al. 2018). These populations are defined as follows:

*Definition 1 (Plausible Population)* A plausible population 𝒫_pop_ generated by direct search methods is a large collection of 𝑁 vectors (or samples) {𝒑_𝟏_, 𝒑_𝟐_, … , 𝒑_𝑵_} ⊂ 𝒫, such that when mapped through a model ℳ: 𝒫 → 𝒪 (generally defined by a system of ordinary differential equations), leads to a distribution of simulated observables {ℳ(𝒑_1_), ℳ(𝒑_2_) … , ℳ(𝒑_𝑁_)} that best approximates the empirical distribution of 𝐷_𝑒𝑥𝑝_ ⊂ 𝒪.

*Definition 2 (Virtual Population)* A virtual population 𝒱_pop_ ⊂ 𝒫_pop_ is a collection of 𝑁^∗^ < 𝑁 vectors (or samples) {𝒑_𝟏_, 𝒑_𝟐_, … , 𝒑_𝑵_∗ }, that minimizes the goodness of fit (GoF) and may also satisfy additional study-specific constraints, such as dropout times (Braniff et al. 2024) or a defined threshold for hepatic fat accumulation in the Van De Pas considered here (Rieger et al. 2022).

### METRICS FOR EVALUATION OF THE PERFOMANCE

To directly compare the performance of DREAM(ZS) with the reference method, we first characterized the Ppop to assess whether it captures sufficient variability in the parameter space. The Ppop was analyzed by measuring the spread of the sampled parameters using violin plots and computing the covariance matrix, as in previous studies [2,9]. Moreover, the performance assessment is structured around the main criteria of diversity, GoF and efficiency (Rieger et al. 2018). A description of these metrics is provided in the following sections.

#### Diversity

To ensure that the Ppop is not only representative of empirical data but also maintains parametric heterogeneity, we assessed diversity using the relative pairwise orientation *d* metric proposed by the reference method (Rieger et al. 2018). This metric quantifies the collinearity between two parameter vectors after scaling and shifting, providing an estimate of how well the Ppop spans the parameter space.

Diversity is assessed by evaluating the empirical cumulative distribution function (ECDF) of all pairwise orientations within the generated population, computed in Eq. 6:

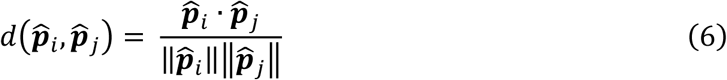

where each scaled and centered parameter vector 𝒑^_𝑘_ is given by:

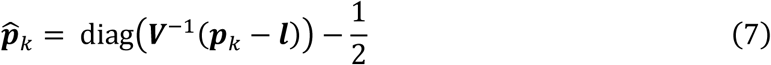

with 𝑽 diagonal of elements Vmm such that 𝑽_𝑚𝑚_ = 𝑢_𝑚_ − 𝑙_𝑚_, representing the difference between the upper 𝑢_𝑚_ and lower 𝑙_𝑚_ bounds of the 𝑚^𝑡ℎ^parameter in the considered population. By definition of the dot product, 𝑑(𝒑^_𝑖_, 𝒑^_𝑗_) corresponds to the angle between parameters vectors with respect to the center of the parameter space. Under ideal conditions where parameters are uniformly distributed across their defined bounds, the ECDF crosses 0.5 at 𝑑 = 0 . On the contrary, shifted ECDFs towards the right or the left indicate a preferred direction of the population in parameter space. Although this metric is deterministic, pairwise computation is computationally expensive for large populations. Therefore, sub-sampling was applied for populations exceeding 1500 individuals.

#### Goodness-of-fit (GoF)

To evaluate the agreement between the simulated population and the observed experimental data, we used, as in the reference method (Rieger et al. 2018), the parametric density estimation (mathematically described in the SM1.4). The Gaussian Multivariate Density Estimation (GMDE), noted as 𝑓^^𝐺^ in SM1.4, measures the similarity between the mean and covariance structure of the simulated and empirical data.

Kolmogorov-Smirnov (KS) test quantifies the discrepancy between the ECDF of the simulated and observed data. Using the one sample KS test, the discrepancy is measured for the 𝑖^𝑡ℎ^observable using the following metric:

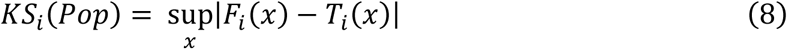

where 𝐹_𝑖_(𝑥) is the ECDF and 𝑇_𝑖_(𝑥) is the corresponding theoretical (target) CDF for observable 𝑖. In our case, 𝐹_𝑖_(𝑥) is obtained by scaling and shifting the 𝑖^𝑡ℎ^observable according to the GMDE to be compared to 𝑇_𝑖_(𝑥), the latter being a standard normal distribution.

The overall GoF metric for the population is computed as:

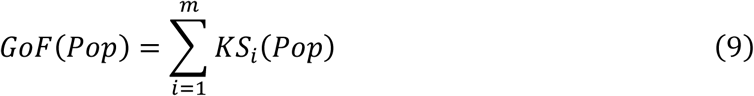

Where 𝑚 is the number of observables, and 𝑃𝑜𝑝 can be either Ppop or Vpop. A lower GoF score indicates a better fit between the virtual population and the experimental data. As noted by Allen et al., a GoF value of 0.2 or less represents an adequate fit (Allen et al. 2016).

#### Efficiency analysis

To compare to the reference method, we evaluated the GoF of the Vpop as a function of the size of Ppop in order to determine the number of PPs required to achieve a given GoF between the generated Vpop and the empirical distribution (𝑓_𝑒𝑥𝑝_). This analysis was extended to assess the stability of the generated Ppop. However, since GoF alone was not sufficient to fully characterize the quality of the simulated population, we complemented the analysis with an evaluation of the mid-point diversity, defined as the ECDF value of diversity at 𝑑 = 0, and analyzed as a function of the Ppop size. Finally, to show the effect of resampling on the Ppop, the expected Vpop size (expressed as a % of the Ppop) was evaluated as a function of the Ppop size.

### CASE STUDY AND DATA

With the objective of using DREAM(ZS) algorithm in pharmacometrics and evaluate its performance in comparison of current methods, the cholesterol metabolism model, originally developed by Van De Pas et al., was taken as a case study (Van De Pas et al. 2012). This model was also used by Allen et al. and Rieger et al. (Allen et al. 2016; Rieger et al. 2018) to illustrate their method. This model consists of a system of nine ordinary differential equations (ODEs) that represent cholesterol production by the liver and its transport through plasma and peripheral tissues. For consistency with previous studies (Allen et al. 2016; Rieger et al. 2018), the model was used to produce steady-state values, which were then compared against empirical cholesterol distributions derived from the National Health and Nutrition Examination Survey (NHANES) dataset (2019). This dataset, representing a diverse US adult population with varying health conditions, provides fasting plasma cholesterol levels for 2942 participants. The joint distribution of high- and low-density lipoprotein cholesterol, (HDL-C) and (LDL-C), respectively, as well as total cholesterol (TC), was well-approximated by a multivariate log-normal distribution, with cholesterol values reported in mg/dL before transformation to log units. The published version of the Van De Pas et al. includes 21 parameters, and the model explicitly calculates HDL-C and non-HDL-C, but not LDL-C or TC. To enable direct comparison with NHANES data, an additional parameter k_22_ was introduced, defined as the ratio of LDL-C to non-HDL-C (Allen et al. 2016).

### SIMULATIONS

All simulations were conducted using MATLAB R2024a, and performed on a i7-13700, 2.10 GHz processor with 64Gb RAM. Each algorithm was run 10 times, providing a sufficient number of repetitions to assess variability and quantify uncertainty. Among these 10 runs, we selected the Ppop with the GoF closest to the mean GoF of the 10 Ppop. On each selected Ppop, 10 Vpop were generated.

## RESULTS

The impact of the algorithm selection on parameter sampling was assessed by comparing the DREAM(ZS) algorithm (C1 and C2) with the MH approach used in the reference method (Rieger et al. 2018) for the generation of Ppop. Moreover, the ability of the algorithms to preserve properties during the transition from plausible to Vpop was also evaluated. In the following, all results for the reference method are based on its original implementation (Rieger 2018). To maintain clarity, the results are organized into two sections: algorithm comparison and properties preservation.

### COMPARISON OF ALGORITHMS

#### Visual comparison of the generated observables

For each of the considered methods, 60000 PPs were generated. Populations generated with DREAM(ZS) initially includes duplicates, which are then filtered to keep only unique samples and be compared to those obtained with the reference method. We recall that this redundancy reflects a deeper exploration of certain regions of the parameter space, representing regions of high probability in the observable space. In the reference method, used as it is for the purpose of comparison, only accepted samples are retained and we did not modify it. To evaluate the ability of DREAM(ZS) to explore the observable space, we compared the multivariate KDE (MKDE) results against those reproduced using the available GitHub source of the reference method (Rieger 2018). The MKDE is used solely to better visualize the difference between the populations. **Fig. 2** illustrates the density estimates distribution for three observable pairs (HDL-C - LDL-C), (HDL-C - TC), and (TC - LDL-C). Here only the results with DREAM(ZS) C1 are included, as similar results were obtained with C2. The main difference is that DREAM(ZS) captures a broader distribution of points, particularly in lower-density regions. This trend is obvious across the comparison of the three observable pairs. The reference method’s approach reinforces the core of the distribution more strongly, leading to a tighter density match in high-probability regions, whereas DREAM(ZS) gives a more distributed density representation, which could be an advantage for capturing a wider range of variability in the observable space.

**Fig. 2.**
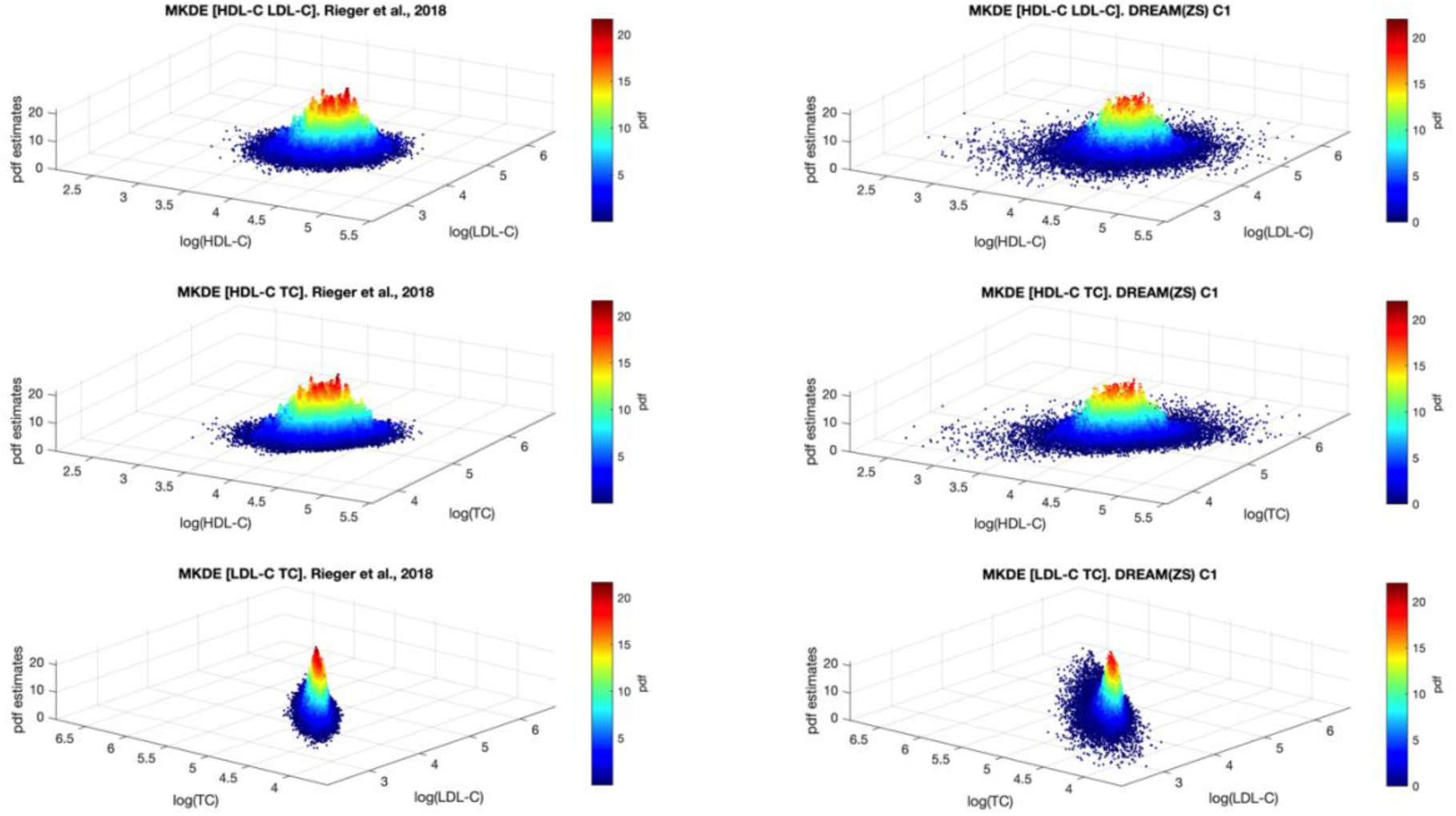
Comparison of multivariate kernel density estimation (MKDE) between the reference methodology (Rieger et al., 2018) and DREAM(ZS) C1. Density estimates for three observable pairs: (HDL-C, LDL-C) (top row), (HDL-C, TC) (middle row), and LDL-C, TC (bottom row). The left column presents MKDE results obtained with Rieger’s source code, while the right column shows results obtained using DREAM(ZS) C1. The color scale represents probability density function (pdf) estimates, with red and blue indicating high-density and low-density, respectively.

#### Parameter space exploration and properties

The exploration of the parameter space across the three methods is summarized in **Fig. 3**, with the violin plots a, i, e for the Ppop. Parameters are normalized with reference to their boundaries between 0 and 1. Results generated using the reference method show accumulations of parameter values near the boundaries, indicating a biased coverage of the parameter space. In contrast, both versions of the DREAM(ZS) algorithm demonstrate a broader exploration, with less accumulation at the boundaries. This highlights the improved sampling efficiency of DREAM(ZS), which exhibits smoother distributions across all parameters. As for the parameter *k_22_*, defined as the ratio between LDL-C and HDL-C, we recall that it was introduced for comparative purposes (Allen et al. 2016) but not explicitly calculated or validated in the cholesterol model (Van De Pas et al. 2012). The boundaries for this parameter were set between 0 and 1; however, the results from the violin plots (**Fig. 3**) suggest that this range may not be realistic, with a mean shifted towards the upper boundary (mean in line with experimental data ratio of 0.85 (Zhang et al. 2016)).

**Fig. 3.**
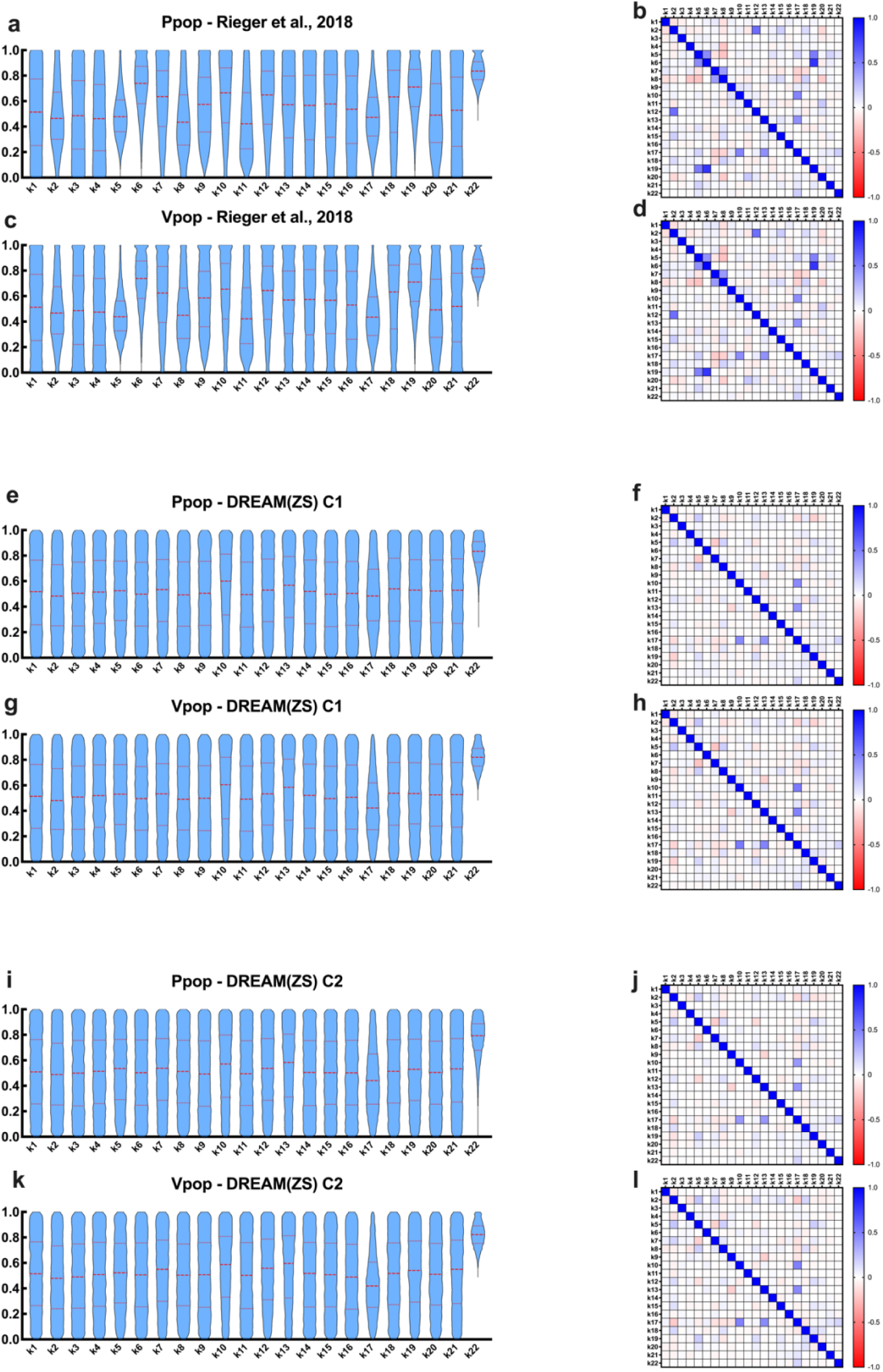
Comparison of parameter space exploration among Rieger et al., 2018 and DREAM(ZS) C1 and C2. Violin plots (left panels) show the distribution of scaled parameter values for Ppop (top) and Vpop (bottom) across different methods: (a, c): Rieger et al., 2018, (e, g): DREAM(ZS) C1, and (i, k): DREAM(ZS) C2. Corresponding correlation matrices (right panels) illustrate parameter relationships: (b, d): Rieger et al., 2018, (f, h): DREAM(ZS) C1, and (j, l): DREAM(ZS) C2. Parameters were scaled using boundary-based scaling to be in the range [0, 1]. Graphs were created using GraphPad Prism version 10.

A clear reduction in parameter correlations is obtained with the DREAM(ZS) method, in contrast with those observed using the reference method (Fig. 3, panels b, f, j). These results highlight the ability of the DREAM(ZS) algorithms to generate more variable and less constrained Ppop due to the underlying sampling strategy. These results are presented for a population size of 60000 for the reference method and ∼27000 for DREAM(ZS) C1/C2. Further analysis on the correlation matrix as a function of Ppop size is provided in the SM1.5 and illustrated in the animation file SM2. Regarding the animation, one sees that the reference method presents strong correlation structures at low Ppop sizes. While those structures do fade away as the size of the population increases, some parameter pairs still stay with high/moderate correlation. For instance (k_5_-k_6_, k_5_- k_19_, k_6_-k_19_, k_10_-k_17_, k_13_-k_17_) are related to various processes critical to cholesterol metabolism, including synthesis, uptake, secretion, and esterification in the liver compartment (Van De Pas et al. 2012). Comparing to DREAM implementations, only few moderate correlated parameters (k_10_- k_17_, k_13_-k_17_) still remain, showing the presence of parameter correlations in the posterior distribution that probably reflects in a more accurate way the underlying model structure and the information content of the data.

To assess the diversity of the Ppop, the cumulative distribution functions (CDFs) of the normalized dot-product metric between pairs of parameter sets defined in Eq. (6) (**Fig. 4**), were computed. Results obtained using the reference method show a rightward shift in the CDF, indicating reduced diversity and a tendency for the Ppop to cluster around similar parameter sets. In contrast, DREAM(ZS) C1 and C2 exhibit CDFs that align more closely with the uniform distribution, demonstrating improved diversity in the sampled parameter space.

**Fig. 4.**
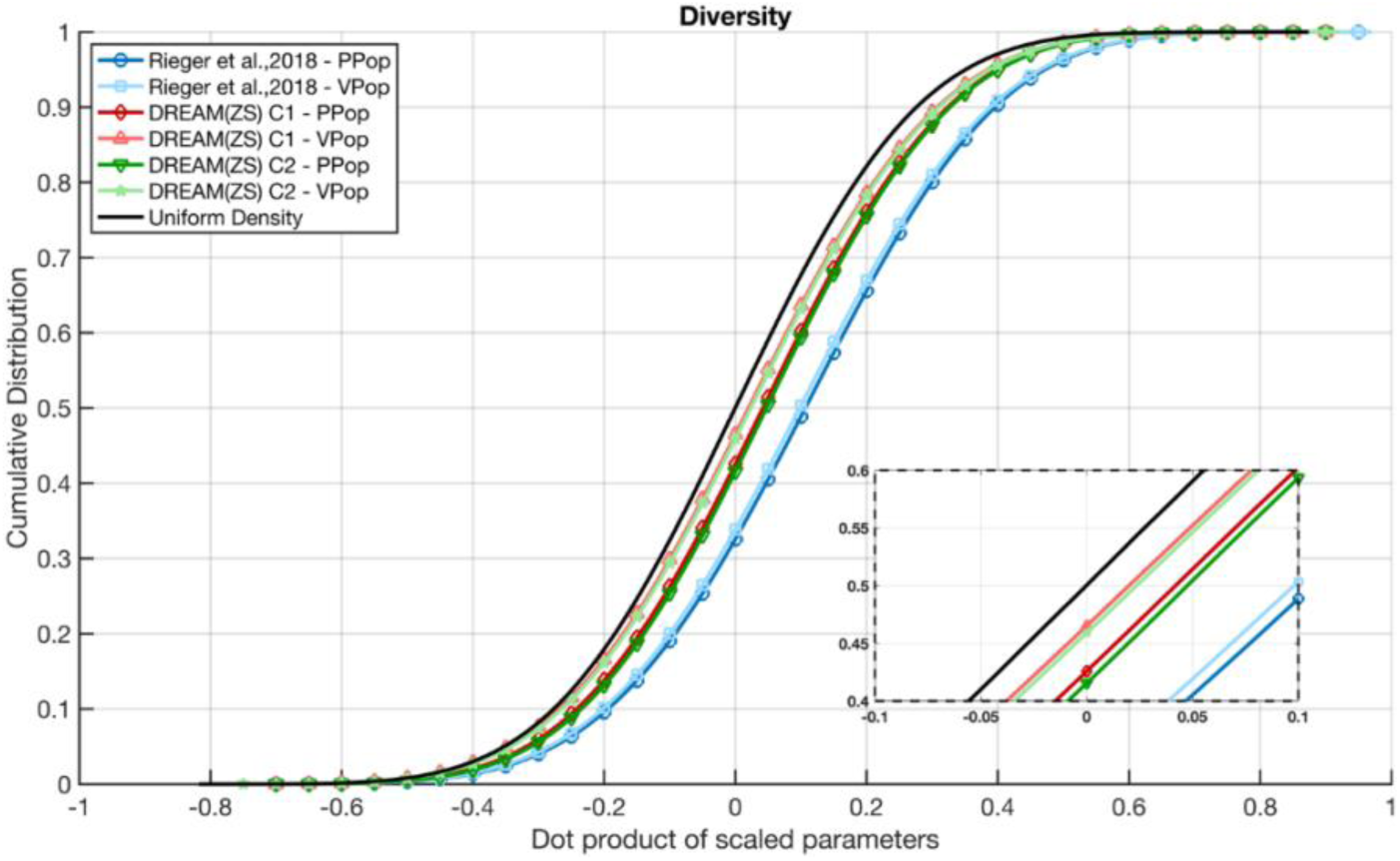
Cumulative distribution functions of scaled parameter dot products comparing diversity across methods. Results obtained with Rieger et al., 2018 source code are in blue (Ppop: circles, Vpop: squares), DREAM(ZS) C1 (red) and DREAM(ZS) C2 (green), shown for both Ppop (diamonds/triangles) and Vpop (lighter shades). The black line represents a uniform density reference. The zoom highlights differences in distribution behavior in the central region.

#### Convergence to the data

The mean GoF was assessed using the Kolmogorov-Smirnov (KS) test for the three observables HDL-C, LDL-C, and TC with a threshold value set at 0.2. The error bars represent the variability across the 10 runs. This GoF has been evaluated for the Ppop with the defined size and for the Vpop after reselection. As shown in **Fig. 5**, the GoF for Ppop obtained with the three methods does not exceed the threshold for each of the single observables, but it does so for the sum of them, with comparable GoF values. The final selection step improves the GoF of Vpop, with a clear reduction in GoF of single observables as well as that of their sum, to values well below the threshold.

**Fig. 5.**
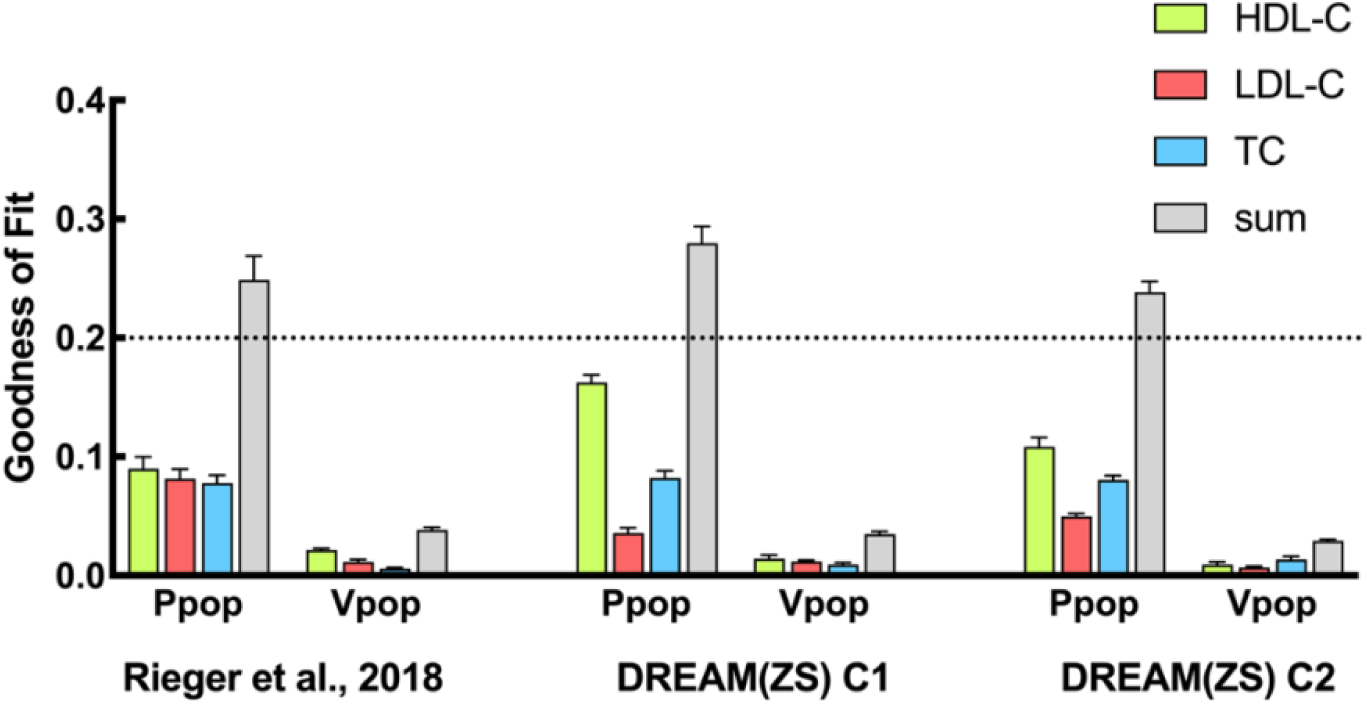
Goodness of Fit (GoF) comparison across methods. Bar plots represent the GoF for different algorithms, comparing Ppop and Vpop across Rieger et al., 2018, DREAM(ZS) C1, and DREAM(ZS) C2. Fit quality is assessed for HDL-C (green), LDL-C (red), and TC (blue), with the sum of all contributions shown in gray. The dotted line serves as a reference threshold value. Error bars indicate variability across the 10 populations generated for each implementation. Graphs were created using GraphPad Prism version 10.

#### Efficiency analysis

The efficiency analysis provides a thorough assessment of both key aspects of plausible and virtual parameter sets: their representativeness of the experimental dataset and their ability to cover the parameter space. Following the reference method’s approach, the GoF of the Vpop was assessed (**Fig. 6a**) to quantify how well the selected Vpop represents the empirical distribution of the data, with lower GoF values indicating a better fit as a function of Ppop size. Each point reflects the average of 10 independently generated VPop, and error bars represent the dispersion. This variability is mostly due to the unstable computation of the 𝛽 factor during the reselection process (see SM1.6). As the size of Ppop increases, the GoF values of all methods evolve to a value below 0.05. However, DREAM(ZS) maintains a more stable and consistently lower GoF throughout, suggesting that the selection of Vpop configurations is more efficient (**Fig.6a**). To evaluate the matching between the generated Ppop and the observed data, GoF of the Ppop was computed for increasing sizes of Ppop (**Fig. 6b**). As Ppop size increases, the GoF of all methods converges towards the same value, with DREAM(ZS) C2 demonstrating a smoother and more gradual decline. The lower GoF values of DREAM(ZS) at smaller population sizes suggest that it requires fewer plausible patients to achieve a well-fitting population. In **Fig. 6c**, the expected mid-point of the diversity ECDF as a function of Ppop size is depicted. We recall that values closer to 0.5 indicate a more diverse parameter space. Diversity was computed using 10 independent samplings per Ppop size; error bars appear only for Ppop sizes ≥1500 due to defined max sample size for computational feasibility. The results show that DREAM(ZS) methods (C1 and C2) maintain a stable diversity mid-point across all Ppop sizes compared to the reference method, which after some fluctuations, stabilizes around 0.35 across population sizes. This suggests that DREAM(ZS) ensures a more isotropic and well-distributed exploration of the parameter space (see SM1.5 for more details). **Fig. 6d** shows the proportion of the total Ppop that qualifies as a Vpop. DREAM(ZS) methods show a more stable Vpop size percentage across Ppop sizes especially starting from Ppop sizes of 1000, suggesting that 1000 may represent an adequate size. As in panel (a), error bars reflect variability over 10 independent replicates per size point. Notably, almost 50% of the Ppop is selected as Vpop in DREAM(ZS) C2, compared to less than 30% in the reference method. This lower eligibility rate questions the quality of the generated Ppop.

**Fig. 6.**
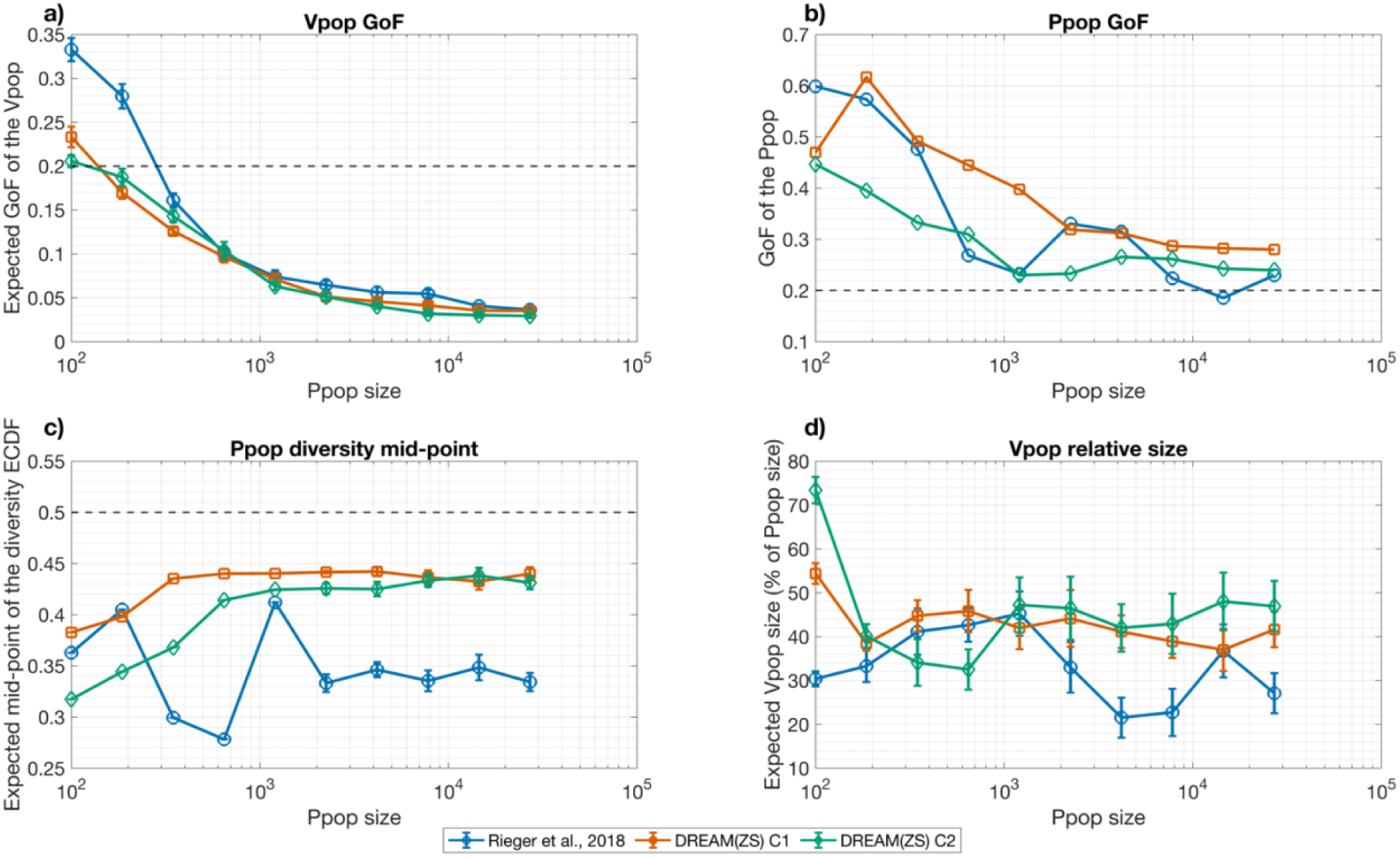
Efficiency analysis across Vpop generation methods. Panels illustrate performance metrics as a function of Ppop size, for three methods: Rieger et al., 2018 (blue), DREAM(ZS) C1 (red), and DREAM(ZS) C2 (green). a) Goodness of Fit (GoF) of the Vpop; b) GoF of the Ppop; c) Diversity of the Ppop, measured as the mid-point of the ECDF of pairwaise cosine distances; d) Relative size of the Vpop (% of the Ppop). Dashed lines indicate reference threshold for GoF (0.2) and diversity (0.5).

Overall, the general stability at different levels appears to be an intrinsic property of DREAM(ZS). Together, this analysis helps determine the optimal population size depending on the application, balancing GoF and diversity in the parameter space.

### PROPERTY CONSERVATION DURING THE TRANSITION FROM PLAUSIBLE TO VIRTUAL POPULATIONS

As raised by Allen et al. (Allen et al. 2016), when selecting a subset of individuals from a larger population, one concern is that the selected subset does not reflect the variability of the original ensemble, which was generated from the biologically plausible range of the parameters. Moreover, the hypothesis inherent to MH (Rieger et al. 2018), and the model itself, may justify an additional step to build Vpop from Ppop. Therefore, the transition from Ppop to VPop was further analysed to determine whether key properties such as parameter exploration, correlation structure and diversity were preserved. As shown in **Fig. 3h** and **3l**, DREAM(ZS) C1 and C2 maintain parameter correlations close to their initial values in Ppop, and parameter distributions in Vpop remain similar to those in Ppop (**Fig. 3e**, **3g**, **3i** and **3k**)

Going from Ppop to Vpop, as ilustrated in **Fig. 4**, an improvement in diversity is obvious for the three methods, as highlighted by the zoomed-in region around a dot product of scaled parameters equal to 0. The diversity of the Vpop generated by the DREAM(ZS) algorithms is closer to the ideal value compared to the reference method. Moreover, the Vpop diversity curves for DREAM(ZS) methods tend to cluster together, instead of being shifted to the right as in the reference method. This confirms that the quality of the initial Ppop directly impacts the resulting Vpop.

## DISCUSSION

Driven by the growing popularity of VPop generation methods and their role in reliably representing physiological, pathological, and pharmacological conditions in patients, we explored the DREAM(ZS) algorithm, which has already proven applicability in the field of hydrology (Laloy and Vrugt 2012). Our objective was to compare this algorithm with state-of-the-art methodology for VPop generation (Rieger et al. 2018), that have gained increasing recognition, using as a case study the cholesterol model developed by Van De Pas (Van De Pas et al. 2012). Considering relevant performance metrics for Vpop algorithms, this study introduces the DREAM(ZS) algorithm to the field of pharmacometrics. Using a well validated model (Van De Pas et al. 2012), we highlight strengths and manifold potential of DREAM(ZS), suggesting that it may be a preferred method not only for its ability to preserve parameter correlations but also to maximize sample diversity within the parameter space, all while matching clinical data distributions.

Compared to current approaches of generation of VPop (Allen et al. 2016; Rieger et al. 2018, 2022; Braniff et al. 2024), DREAM(ZS) allows for a better exploration of parameter space, thanks to the use of differential evolution techniques. Moreover, DREAM(ZS) has shown its ability in capturing lower-density regions in the observable space (**Fig. 2**), broadening the coverage of the core distribution; this can be advantageous for modeling rare clinical phenotypes, providing an alternative to traditional methods (Allen et al. 2016; Rieger et al. 2018; Sinisi et al. 2020). Parameter distributions were also more evenly spread, with fewer samples collapsing against bounds compared to the reference method, as illustrated in **Fig. 3**. These differences suggest that DREAM(ZS)’s exploration strategy better avoids over-sampling of parameter extremes, handling boundary conditions by using reflective or adaptive strategies, which help preserve a better exploration of the parameter space. In contrast, the reference method uses boundary truncation to limit parameters, a strategy that, as discussed by Vrugt, can make the samples less varied, cause too many values to accumulate at the edges, and lead the algorithm to get stuck instead of properly exploring the full space (Vrugt 2016).

One of the primary strengths of DREAM (ZS) is its ability to handle complex distributions and achieve efficient convergence. DREAM(ZS) excels at exploring multimodal parameter spaces, ensuring a very good coverage of possible populations (Vrugt et al. 2009; Vrugt 2016).

While the internal mechanisms of DREAM(ZS) are quite sophisticated, its practical application remains user-friendly. The algorithm is designed to be accessible for researchers, facilitating its integration into various modeling workflows without requiring extensive computational expertise. Moreover, by enabling the creation of diverse, representative Ppop, the algorithm may be used for drug development and trial simulations. This, in turn, allows for better optimization of trial designs, potentially improving the success of therapeutic outcomes.

While no extensive analysis was performed here to measure the difference in computation time, DREAM(ZS) seems to run faster than the reference method. Other studies, using machine learning-based surrogate models, are actively exploring this aspect with the primary goal of accelerating VPop generation (Myers et al. 2023; Iwata and Saito 2025).

The established use and versatility of the DREAM algorithm family is well-documented in the literature and supported by a range of software tools, including the CroptimizR R package for agricultural model calibration (Buis et al. 2023) and the original MATLAB implementation (Vrugt 2025). DREAM(ZS) has been successfully applied to a wide variety of problems, including sampling from complex target distributions, inferring parameters in time series models, and performing diagnostic evaluations of environmental and systems models (Vrugt 2016). The associated MATLAB toolbox includes over 30 ready-to-run examples that can be adapted to diverse modeling tasks, such as ODE-based systems, stochastic simulations, and empirical data fitting. This makes DREAM(ZS) particularly applicable for broader QSP applications, including the generation of VP across disease areas. Furthermore, advanced algorithms of the DREAM family have been developed for high-dimensional parameter spaces (more than 200 parameters), making them potentially suitable for large-scale models (Vrugt 2016).

The use of DREAM(ZS) in the context of the cholesterol metabolism with steady-state outputs was intended to allow comparison with current methods. Because of the above-mentioned capacities of this algorithm, one can expect that DREAM(ZS) will reveal its full potential when applied to more complex models and time variant outputs. Investigating additional performance metrics will also be crucial for refining the algorithm and adapt it to pharmacometrics area.

Our analysis points towards a deeper examination of the proposed metrics in previous studies (Allen et al. 2016; Rieger et al. 2018). As defined in Eq. (6), a given diversity value does not fully inform the exploration of the parameter space. Indeed, it is possible to construct a set of different Vpop (differing in their exploration qualities) which exhibits the same diversity ECDF. It is also worth mentioning that diversity is not fully related to the correlation between parameters. While its shape (of the diversity curve) is influenced by the dimensionality of the problem (22 parameters here), it is also linked to the size of the considered population (Fig.S1). However, a stability in this change of the shape has been noticed for DREAM(ZS) compared to the reference method, as shown in the SM1.4. Future work could explore diversity issues using other methodologies. As for the GoF metric, it relies solely on the marginal distributions of each observable. This ensures accurate representation of univariate data distributions but neglects their combined distributions. Furthermore, alternative strategies should be investigated to minimize resampling steps or optimize selection criteria without requiring further adjustments as through the use of 𝛽 optimization to go from the Ppop to the Vpop. Indeed, as shown in a recent work by Braniff et al. (see Fig. 3), the 𝛽 optimization step was not involved (Braniff et al. 2024).

Another important consideration is the selection of chain elements after the burn-in period, which is inherent to DREAM(ZS) implementation. Initial samples may not yet represent the desired distribution, and therefore burn-in period should be considered in MH.

The criteria for determining the optimal size for a Vpop carry implications for future clinical applications. While large Vpop are often used, our results suggest that a smaller, well-characterized subset, selected based on good properties of diversity and GoF, can be sufficient. In perspective, this approach highlights the potential of minimal size of Vpop to detect meaningful treatment differences, which is relevant to inform clinical trials design (Braniff et al. 2024).

As a next step, clustering algorithms could be applied to the Vpop generated using DREAM(ZS). Since this algorithm produces a diverse set of parameter combinations, clustering can help identify groups of VP that share similar biological characteristics or model outputs. For example, by first reducing the dimensionality of the parameter space using UMAP, and then applying methods like k-nearest neighbors (kNN), it becomes possible to detect subgroups with similar risk profiles or treatment responses. This type of analysis could improve interpretability, reveal hidden structure within the VPop, and support more targeted clinical trial simulations for instance, by simulating outcomes separately for different patient subtypes or prioritizing biomarkers within each cluster.

## CONCLUSION

When it comes to computational aspects, the biological field can largely benefit from methods developed in completely different areas. We here investigated an algorithm introduced and used in hydrology, but whose characteristics can be adapted to QSP and VPop generation. With the aim of providing an interesting alternative to what already exists for Vpop generation in pharmacology, we described its practical use, analysed its strengths as well as limitations, and provided a ready-to-be-used code for those who are interested to readily apply it to their own problems, offering potential added value for the QSP community.

## Acknowledgments

Support was provided by NSERC (Collaborative Research and Development Grants), in partnership with Syneos Health and Pfizer; FRQNT-Projet d’équipe, and from Prompt. MS received financial support by the Fonds de recherche du Quebec – Nature et technologies (FRQNT) and the Centre Interdisciplinaire de Recherche sur le Cerveau et l’Apprentissage (CIRCA).

## Author contributions

MS and TB contributed equally to this work. Conceptualization: MS, TB, DZ, POT, FN; Methodology: MS, TB, DZ, FN; Code development and simulations: TB, MS, MP, DZ; Investigation: TB, MS, MP, DZ; Funding acquisition: FN, POT; Writing – original draft: MS; Writing – review & editing: MS, TB, DZ, POT, FN; Supervision: FN, DZ.

All authors reviewed and approved the final manuscript.

## Ethics approval

Not applicable.

## Data availability

The code necessary to reproduce the results of this study has been made available to reviewers during the submission phase. Upon acceptance for publication, a public GitHub repository will be created containing the full set of scripts, data and documentation. A DOI and repository link will be provided in the final published version of the paper.

