## Supplementary material for "Generation of virtual populations for quantitative systems pharmacology through advanced sampling methods": Supplemenary Material 1

#### Supplementary Material (SM1)

Miriam Schirru[1], Tristan Brier[1], Maxime Petit[1], Didier Zugaj[2], Pierre-Olivier Tremblay[2], and Fahima Nekka[1,3]

[1]Laboratoire de recherche en pharmacométrie, Faculté de pharmacie, Université de Montréal, Montréal, Canada
[2]Syneos Health, Clinical Pharmacology, Québec, Canada
[3]Centre de recherches mathématiques, Université de Montréal, Montréal, Canada

**SM1.5 Efficiency Analysis: Diversity** **7**

**SM1.6 Stability of the scaling factor $\beta$ for Vpop selection** **9**

### List of Algorithms

1 $\mathcal{L}_{\mathrm{C1}}$ Computation Pseudocode . . . . . 4
2 $\mathcal{L}_{\mathrm{C2}}$ Computation Pseudocode . . . . . 4
3 GoF as a function of $\beta$ (Score function for SA) . . . . . 9

#### SM1.1 Bias corrected likelihood $\mathcal{L}_{\mathbf{C2}}$

##### SM1.1.1 Derivation using Bayes' theorem

The generation of the Vpop can be approached as a sampling problem: we aim to construct a sampling of $\mathcal{P}$ (the parameter space) such that the distribution of its outputs in $\mathcal{O}$ (the observable space) matches the distribution of data. Following the methodology of MCMC MH algorithms, we then aim to construct a Markov chain sampling the distribution of the parameter vectors $p \in \mathcal{P}$ given the model mapping $\mathcal{M}$ and the data $D$.

The desired posterior distribution of $\mathbf{p}$ (a random vector whose values are the parameter vectors), $f\left(\mathbf{p} = p \mid \mathcal{M} \circ \mathbf{p} = \mathcal{M}(p), \mathbf{D} = D\right)$, is derived using Bayes' theorem. It combines the likelihood $f\left(\mathcal{M}(p), D \mid p\right)$ of the data and model outputs given $p$, the prior distribution of $\mathbf{p}$, $f(p)$, and the marginal likelihood $f(\mathcal{M}(p), D)$ of the model outputs and data as follows:

$$f\left(\mathbf{p} = p \mid \mathcal{M} \circ \mathbf{p} = \mathcal{M}(p), \mathbf{D} = D\right) = \frac{f(p)\, f\left(\mathcal{M}(p), D \mid p\right)}{f(\mathcal{M}(p), D)} \tag{SM1.1.1}$$

Considering $\mathbf{D}$ is independent of both $\mathbf{p}$ and $\mathcal{M} \circ \mathbf{p}$, the posterior can be simplified and viewed as a weighing function of the model outputs $\mathcal{M}(p)$ according to their relevant input $p$ and data $D$:

$$f\left(\mathbf{p} = p \mid \mathcal{M} \circ \mathbf{p} = \mathcal{M}(p), \mathbf{D} = D\right) = \frac{f(p)\, f\left(\mathcal{M}(p) \mid p, D\right)}{f(\mathcal{M}(p))} \tag{SM1.1.2}$$

To derive SM1.1.2, knowing the conditional probability density is expressed by $f\left(x \mid y\right) = \frac{f(x,y)}{f(y)}$, the likelihood $f\left(\mathcal{M}(p), D \mid p\right)$ becomes:

$$\begin{aligned} f\left(\mathcal{M}(p), D \mid p\right) &= \frac{f((\mathcal{M}(p), D), p)}{f(p)} \\ &= \frac{f(\mathcal{M}(p), (p, D))}{f(p)} \\ &= \frac{f(D, p)\, f\left(\mathcal{M}(p) \mid D, p\right)}{f(p)} \end{aligned}$$

Reintroducing this last form in Eq.(SM1.1.1) gives:

$$\begin{aligned} f\left(\mathbf{p} = p \mid \mathcal{M} \circ \mathbf{p} = \mathcal{M}(p), \mathbf{D} = D\right) &= \frac{f(p)}{f(\mathcal{M}(p)), D} \frac{f(D, p)\, f\left(\mathcal{M}(p) \mid D, p\right)}{f(p)} \\ &= \frac{f(D, p)\, f\left(\mathcal{M}(p) \mid D, p\right)}{f(\mathcal{M}(p), D)} \end{aligned}$$

Considering finally that $\mathbf{D}$ is independant of both $\mathbf{p}$ and $\mathcal{M} \circ \mathbf{p}$, one obtain:

$$f\left(\mathbf{p}=p \mid \mathcal{M}\circ\mathbf{p}=\mathcal{M}\left(p\right),\mathbf{D}=D\right)=\frac{f\left(D\right)f\left(p\right)f\left(\mathcal{M}\left(p\right)\ \middle|\ D,p\right)}{f\left(\mathcal{M}\left(p\right)\right)f\left(D\right)}$$

$$f\left(\mathbf{p}=p \mid \mathcal{M}\circ\mathbf{p}=\mathcal{M}\left(p\right),\mathbf{D}=D\right)=\frac{f\left(p\right)f\left(\mathcal{M}\left(p\right)\ \middle|\ D,p\right)}{f\left(\mathcal{M}\left(p\right)\right)} \quad \text{(SM1.1.3)}$$

This revised target distribution for the parameters integrates now a non-constant normalization factor, namely $f\left(\mathcal{M}\left(p\right)\right)$. To include the computation of this factor in DREAM(ZS), we redefine the likelihood as $\mathcal{L}_{\mathrm{C2}}$, with

$$\mathcal{L}_{\mathrm{C2}}\left(\mathcal{M}\left(p\right)\ \middle|\ D,p\right)=\frac{f\left(\mathcal{M}\left(p\right)\ \middle|\ D,p\right)}{f\left(\mathcal{M}\left(p\right)\right)} \quad \text{(SM1.1.4)}$$

The following representation are used in the main document:

- $f\left(p\right)$ is $f_{prior}\left(p\right)$
- $f\left(\mathbf{p}=p \mid \mathcal{M}\circ\mathbf{p}=\mathcal{M}\left(p\right),\mathbf{D}=D\right)$ is $f_{post}\left(p \mid \mathcal{M}\left(p\right)\right)$
- $f\left(\mathcal{M}\left(p\right)\ \middle|\ D,p\right)$ is given by the experimental density $f_{exp}\left(\mathcal{M}\left(p\right)\right)$
- $f\left(\mathcal{M}\left(p\right)\right)$ is the density of the pushforward prior by $\mathcal{M}$, $f_{sim}\left(\mathcal{M}\left(p\right)\right)$

##### SM1.1.2 Construction of the density $f\left(\mathcal{M}\left(p\right)\right)$

The denominator in Eq.(SM1.1.2) corresponds to the probability density of the model outputs under the prior distribution of parameters. Unlike many standard applications of Bayes' theorem, this quantity cannot be assumed constant and must therefore be explicitly computed. However, $\mathcal{M}$ may lack analyticial properties such as continuity and invertibility to explicitly construct this pdf as a function of the prior.

To circumvent this complication, we resort to an empirical estimation of $f\left(\mathcal{M}\left(p\right)\right)$ as follows:

1. Generate a sample $\{\mathbf{p}_i\}$ of sufficient size using the prior of $\mathbf{p}$. In this study, the size was chosen to match the number of data points. LHS sampling was used since the prior is a uniform distribution. For more complex priors, DREAM(ZS) itself could be used to perform the sample.

2. Apply the model on the sample to get the set $\{\mathcal{M}\left(\mathbf{p}_i\right)\}$. This sample follows the pushforward pdf $f\left(\mathcal{M}\left(p\right)\right)$.

3. Construct an estimator of the pdf $f\left(\mathcal{M}\circ\mathbf{p}\right)$ using $\{\mathcal{M}\left(\mathbf{p}_i\right)\}$ with a kernel density estimator (Eq.SM1.4.3), denoted $\hat{f}\left(\mathcal{M}\circ\mathbf{p}\right)$.

With the estimator $\hat{f}\left(\mathcal{M}\circ\mathbf{p}\right)$ in hand, the denominator of SM1.1.2 is then evaluable as a function of $p$.

#### SM1.2 Likelihood Computation Pseudocode

The likelihood function is a central component of the method and must be implemented in a modular manner. In particular, the function should accept a single input (the parameter vector `p`, representing $p$ in the pseudocode) and return the corresponding likelihood or log-likelihood (see [4] for details). All necessary computations to evaluate the likelihood from `p` must be performed internally by the function, without relying on additioonnal arguments.

Algorithms 1 and 2 outline the computation steps for the $\mathcal{L}_{C1}$ and $\mathcal{L}_{C2}$ likelihood formulations, respectively

**Algorithm 1** $\mathcal{L}_{\mathrm{C1}}$ Computation Pseudocode

1. Construct the experimental density $\hat{f}_{\mathrm{exp}}$ as a fixed object callable from within the function (f.e.: `persistent` command in MatLab [3])
2. Compute observables $\mathtt{Obs} = \mathcal{M}(\mathtt{p})$. This task can be relegated to an external function script called within the likelihood function script.
3. Evaluate and return $\hat{f}_{\mathrm{exp}}(\mathtt{Obs})$

**Algorithm 2** $\mathcal{L}_{\mathrm{C2}}$ Computation Pseudocode

1. Construct the experimental density $\hat{f}_{\mathrm{exp}}$ as a fixed object callable from within the function (f.e.: `persistent` command in MatLab [3])
2. Construct the pushed-forward prior $\hat{f}_{\mathrm{sim}}$ as a fixed object callable from within the function using the method detailed in sec. SM1.1.2.
3. Compute observables $\mathtt{Obs} = \mathcal{M}(\mathtt{p})$. This task can be relegated to an external function script called within the likelihood function script.
4. Evaluate and return $\hat{f}_{\mathrm{exp}}(\mathtt{Obs})/\hat{f}_{\mathrm{sim}}(\mathtt{Obs})$

In addition, the computed observables can be saved from within the likelihood script to avoid recomputation after the Ppop generation.

#### SM1.3 DREAM Diagnostic Tools

##### SM1.3.1 Overview of used diagnostic to

The DREAM matlab toolbox provides many diagnostic and visualisation tools, all of which are detailed in [4]. We describe here the ones useful for our work:

- **R-diagnostic $\hat{R}$:** this metric measures the convergence of the chains to the target distribution. When the multivariate $\hat{R}$ drops below a threshold value of 1.2, the chains are considered to have converged to the target distribution. The iteration at which this threshold is reached defines the burn-in period. All samples prior to this point are discarded.

- **Acceptance rate (AR):** the AR corresponds to the proportion of proposed parameter sets that are accepted. As mentionned in the DREAM manual [4], the AR should range between 15% and 30%. Deviations from these values indicate sampling inefficiencies: a high AR may suggest poor exploration due to proposals being too similar, while a low AR may reflect for example an inadequate autocorrelation between samples, both slowing the convergence of the chains or providing wrongful results. The AR can typically be tuned by modifying the MH internal parameters. In this study, the AR also approximates the proportion of unique PPs, as each accepted parameter set corresponds to a distinct PP.

- **Autocorrelation:** For each parameters, the autocorrelation function measures the correlation between 2 samples which are some `Lag` time appart. The speed at which those curves decay indicates the quality of mixing between chains. We use it mostly to make sure the chains are mixing well enough during the process and to heuristically interpret the intervariability of the sampling, a sought after quality of a Vpop.

- **Chain Trajectories:** Visualising the chain trajectories is most useful to assess exploration of the parameters space. We can understand more clearly if there are some possible problems during the proposition mechanism or if the chains get stuck in some region of the parameters space.

##### SM1.3.2 Estimating the number of iterations yielding a target Ppop size

Using the given diagnostic tools, we assess the target number of iterations as follows:

1. Set some target number of PPs in the Ppop, say 20 000 for example.

2. With small number of iterations, run DREAM(ZS) successivly to assess the burnin time and the approximate acceptance rate post-burnin. Say for example the burnin is 2500 and the AR is 30%.

3. The number of iterations is then set as $T = \frac{\#\text{PPs}}{\text{AR}*N} + \text{burnin}$ with $N$ the number of chains. If we run 3 chains in this example, $T = \frac{20000}{0.3*3} + 2500 \approx 24730$.

This method ensures that when the burnin phase is removed and the duplicates filtered out, around 20000 PPs will remain.

#### SM1.4 Density Estimators

##### SM1.4.1 Parametric Estimation (Multivariate Gaussian)

Multivariate gaussian density estimator. Given some data set $D = \{\mathbf{d}_i\}$ of size $n$, we can compute the experimental mean vector $\boldsymbol{\mu}_{\text{exp}}$ and covariance matrix $\boldsymbol{\Sigma}_{\text{exp}}$ given by:

$$\boldsymbol{\mu}_{\text{exp}} = \frac{1}{n}\sum_{i=1}^{n} \mathbf{d}_i \quad , \quad \boldsymbol{\Sigma}_{\text{exp}} = \frac{1}{n-1}\sum_{i=1}^{n} \left(\mathbf{d}_i - \boldsymbol{\mu}_{\text{exp}}\right)\left(\mathbf{d}_i - \boldsymbol{\mu}_{\text{exp}}\right)^T \tag{SM1.4.1}$$

From the mean and covariance matrix, the gaussian density estimator is:

$$\hat{f}^{G}_{\text{exp}}(\mathbf{x}) = \frac{1}{(2\pi)^{m/2}\left|\boldsymbol{\Sigma}_{\text{exp}}\right|^{1/2}} \exp\left(-\frac{1}{2}\left(\mathbf{x} - \boldsymbol{\mu}_{\text{exp}}\right)^T \boldsymbol{\Sigma}^{-1}_{\text{exp}}\left(\mathbf{x} - \boldsymbol{\mu}_{\text{exp}}\right)\right) \tag{SM1.4.2}$$

##### SM1.4.2 Non-Parametric Estimation (Kernel Density Estimation)

Instead on relying on a specific distribution to model the target, we could use the following non-parameteric estimator.Given a kernel $K(\cdot)$ (e.g., Gaussian kernel) and a bandwidth matrix $H$, the kernel-based density estimation is:

$$\hat{f}^{K}_{\text{exp}}(\mathbf{x}) = \frac{1}{n}\sum_{i=1}^{n} \frac{1}{\sqrt{H}} K\left(H^{-1/2}\left(\mathbf{x} - \mathbf{d}_i\right)\right) \quad , \quad \text{where } K(\mathbf{u}) = \frac{1}{(2\pi)^{m/2}} \exp\left(-\frac{1}{2}\left|\left|\mathbf{u}\right|\right|^2\right) \tag{SM1.4.3}$$

#### SM1.5 Efficiency Analysis: Diversity

During the efficiency analysis, the evolution of the mid-point diversity is assessed in relation to the Ppop size. This section presents the full diversity ECDF for each implementation in relation to the Ppop size.

Figure S1 presents the ECDF of diversity for each implementation. The most striking observation concerns convergence: DREAM C1/ C2 show a consistent progression toward a limiting diversity ECDF, whereas the reference method (Rieger et al., [2]) shows a more irregular and less stable trajectory. This suggests that DREAM(ZS) performs a more consistent and isotropic exploration of the parameter space from the outset of the sampling process.

In the reference method, the ECDFs display differences in curvature, which, as discussed in the main text, are linked to the effective dimensionality of the sample space. Supporting this, the supplemental animation `Correlation_Matrices_Evolution.mp4` illustrates that the reference method produces strong inter-parameter correlations at small Ppop sizes. In contrast, both DREAM(ZS) implementations achieve rapid decorrelation. Since the correlation structure reflects the degrees of freedom in the population, these differences help explain the contrasting ECDF shapes and further highlight DREAM's superior diversity performance.

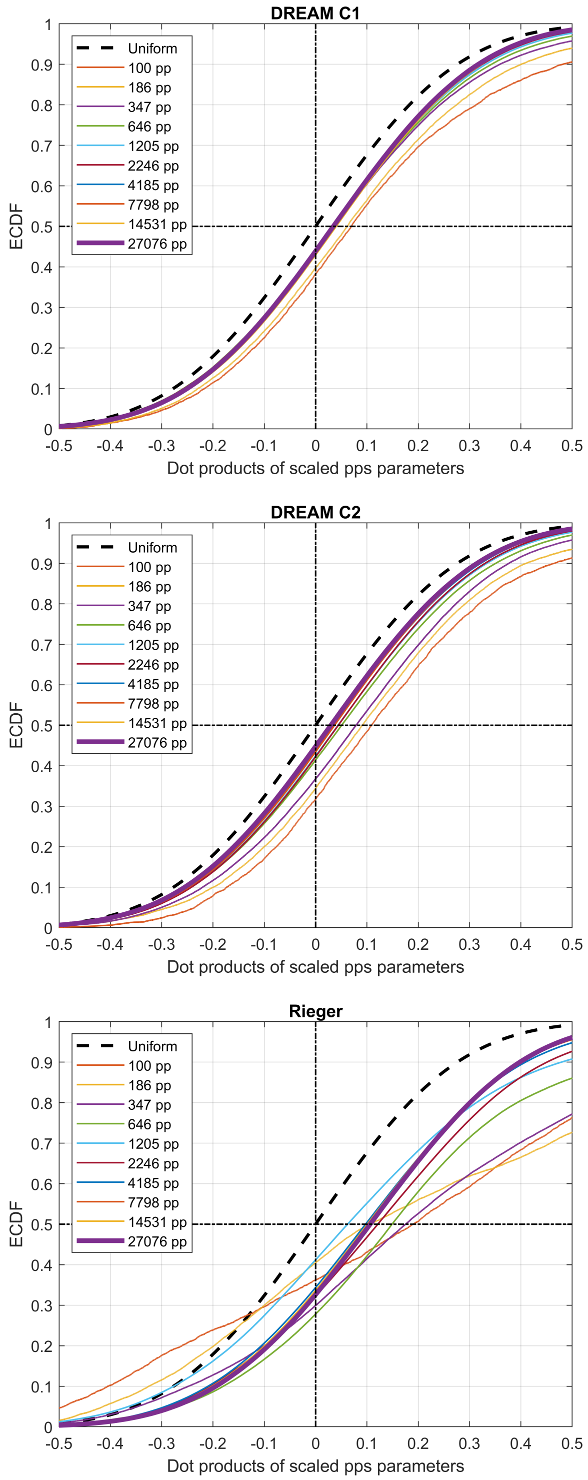

Figure S1: Diversity ECDF evolution as a function of the Ppop size for DREAM(ZS) (top C1 - middle C2) and Rieger (bottom). DREAM exhibits consistent convergence toward a stable limiting ECDF, whereas Rieger's method shows more irregular behavior across iterations

#### SM1.6 Stability of the scaling factor $\beta$ for Vpop selection

Following the methodology of Allen et al.[1], each patient $p_j \in P_{\text{pop}}$ is assigned a probability for being included in the $V_{\text{pop}} \subset P_{\text{pop}}$:

$$\Pr\left( S\left(p_j\right) = 1 \;\middle|\; \mathcal{M}\left(p_j\right) = r \right) \approx \beta \frac{f_D\left(r\right)}{\rho_{\text{pl}}\left(r\right)} \quad , \quad S\left(p_j\right) = \begin{cases} 1 & \text{, if } p_j \in V_{\text{pop}} \\ 0 & \text{, otherwise} \end{cases} \tag{SM1.6.1}$$

where $f_D$ is the data probability density estimator and $\rho_{\text{pl}}$ is density estimator of the Ppop, both evaluated at $r \in O$. This probability of inclusion relies on the free parameter $\beta$, which acts as a scaling factor. The estimation of this scaling factor is operated through an optimization process (simulated annealing, SA) with the goal of minimizing the GoF of the selected Vpop.

However, the optimization problem is not mathematically well defined since the score function for the process is itself stochastic. Algorithm 3 presents a pseudocode outlying the computation of the score function.

**Algorithm 3** GoF as a function of $\beta$ (Score function for SA)

1. Set value for number of runs, $n$. Initialize GoF container $K$

2.:

**for** $1 \leq j \leq n$ **do**

  i) Perform selection according to Eq.SM1.6.1

  ii) Compute GoF as sum of ks-tests

  iii) Save GoF in $K\left(j\right)$

**end for**

3. **return** mean($K$)

The stochastic nature of the score function specifically appears in step 2.i), where the selection is random. This randomness has a significant impact: as shown in Figure S2, it introduces artificial jagged fluctuations in the score curve, leading to numerous local minima. These local minima can mislead the optimization algorithm into prematurely identifying suboptimal solutions as the global minimum. Furthermore, beyond these fluctuations, the lower panel of Figure S2 illustrates that the cost function remains nearly flat around the region corresponding to the global minimum. Together, these two effects result in high variability in the estimated value of $\beta$.

This variability in $\beta$, in turn, translates to substantial fluctuations in the final size of the selected VPop after the acceptance-rejection step. While the Vpop size remains stable for any fixed $\beta$, Figure S3 demonstrates that the $\beta$ values selected by the optimization procedure fall within a steeply increasing region of the Vpop size curve, with Vpops ranging from 8400 to 15500 individuals.

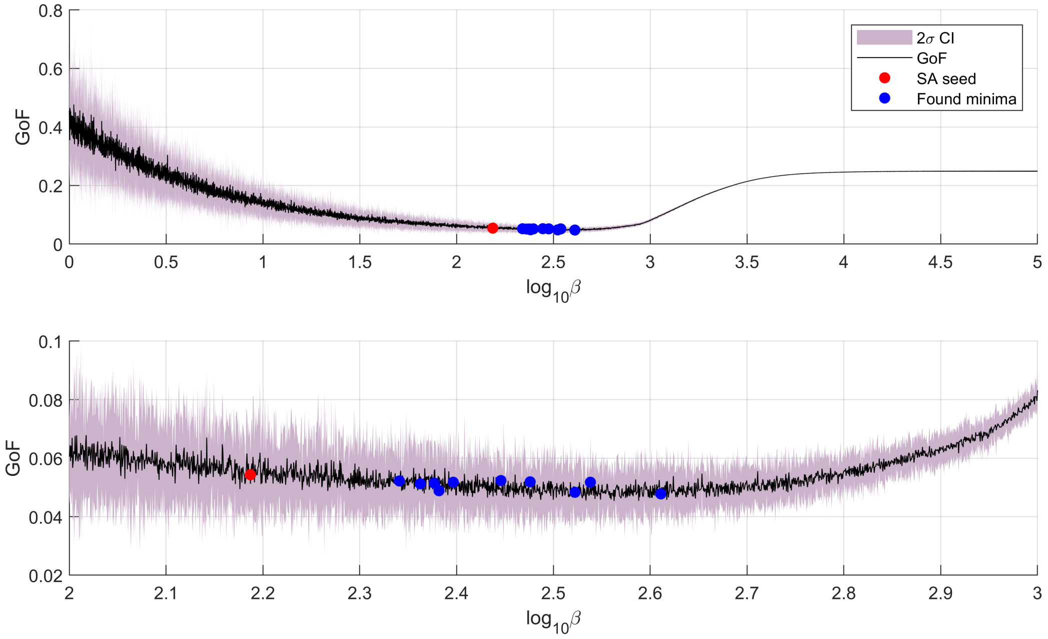

Figure S2: Variation of the GoF as a function of $\beta$ with 95% CI. The plot illustrates the stochastic nature of the score function, with high variability at low $\beta$ values and jagged fluctuations across the curve. These effects are evident in the zoomed-in region.

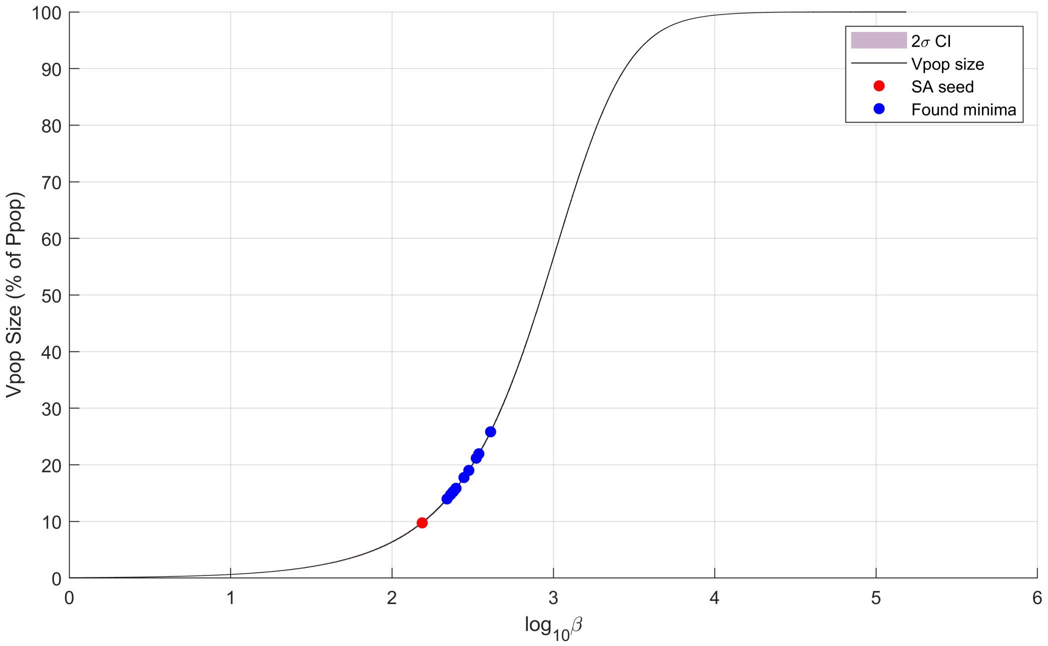

Figure S3: Variation of the Vpop size as a function of $\beta$ with 95% CI. This plot demonstrates the stability of the Vpop size with respect to a $\beta$, as reflected by the negligible width of the CIs.
